# An Alport variant illuminates the bioactivity of the collagen IV ^α565-α121^ scaffold in Bowman’s capsule

**DOI:** 10.64898/2026.02.23.707599

**Authors:** E Pokidysheva, R. Koirala, B. Clarke, E. Delpire, S. Boudko, BG. Hudson

## Abstract

Alport syndrome (AS) is a major cause of chronic kidney failure and affects millions of people worldwide. Pathogenic variants in COL4A3, COL4A4, and COL4A5, which encode the collagen IV ^**α345**^ scaffold, compromise glomerular basement membrane (GBM) structure and function. However, the molecular mechanisms linking the >5,000 reported variants to disease pathology remain poorly understood. To address this gap, we previously examined a distinctive variant, an 8-amino acid “Z-appendage”, added to the NC1 domain of the α3 chain. Knock-in mice carrying this variant developed GBM abnormalities and proteinuria, implicating the NC1 hexamer as a critical determinant of GBM function and suggesting that the hexamer surface contains bioactive sites that may mediate signaling and/or organization of macromolecular complexes. Given that approximately 80% of AS cases are associated with COL4A5 variants, including many within the NC1 hexamer, we asked whether relocating the Z-appendage from the α3 NC1 subunit to the α5 subunit produces similar pathology. Strikingly, *Col4a5-Z* mice did not develop proteinuria and showed only minor changes in GBM morphology. In contrast, the variant induced marked thickening of Bowman’s capsule, accompanied by increased deposition of the collagen IV ^**α121**^ scaffold, increased fibrillar collagen, and cellular deposits. Structural modeling predicts that the collagen IV ^**α565–α121**^ scaffold bearing two Z-appendages adopts an aberrant secondary structure that may stiffen the scaffold and occlude binding sites. Together, these findings reveal a bioactive role for the collagen IV ^**α565–α121**^ scaffold in the Bowman’s capsule basement membrane, with potential implications for other α565–α121containing tissues such as the aorta and bladder.

## Introduction

Collagen IV is a critical structural and functional component of the glomerular basement membrane (GBM), a key element of the renal filtration barrier. In mammals, collagen IV consists of six distinct chains (α1–α6), which assemble into three major isoforms: α121, α345, and α565. The α345 isoform predominates in the GBM, where it provides essential mechanical strength and filtration efficiency [1], [2], [3].

Alport syndrome (AS), the most common hereditary glomerular disorder, arises from dysfunction in the collagen IV^**α345**^ network due to mutations in the *COL4A3, COL4A4*, or *COL4A5* genes [4]. These mutations impair the proper assembly of collagen IV, leading to either a complete absence of the α345 scaffold in the GBM or deposition of a defective scaffold [5]. Approximately 85% of AS cases are caused by mutations in the *COL4A5* gene, which results in an X-linked form of the disease, while the remaining 15% are due to autosomal recessive mutations in *COL4A3* or *COL4A4* [4], [6]. Clinically, AS is characterized by hematuria, proteinuria, and progressive renal failure [7], [8], [9]. The severity of the disease varies depending on the specific mutation, with some individuals experiencing milder forms or incomplete penetrance, complicating diagnosis and genetic counseling [5], [10], [11].

Research indicates that individuals with COL4A5 mutations who retain a partially functional α345 network in the GBM experience a slower progression to kidney failure compared to those who completely lack the network. The specific mutation type plays a pivotal role in determining the severity and progression of the disease [5], [11]. This highlights the essential role of the collagen IV^**α345**^ assembly in preserving kidney function. Another α5-containing collagen IV scaffold, α565, is predominantly found in Bowman’s capsule and is also affected in cases of X-linked Alport syndrome [12], [13], [14]. However, the mechanisms through which defective collagen IV scaffolds contribute to renal pathology remain largely unknown.

Among the various pathogenic variants in collagen IV genes, Pokidysheva et al. identified one associated with familial Goodpasture disease, which results in an 8-amino acid peptide extension at the C-terminal NC1 domain of the α3 chain [15]. A knock-in mouse model incorporating this peptide—referred to as the Zurich (Z-) appendage—exhibited moderate proteinuria, glomerulosclerosis, and GBM abnormalities resembling Alport syndrome. Given that approximately 85% of AS cases are linked to mutations in the α5 chain [16], and multiple pathogenic variants are found withing the NC1 domain [17], we used the Z-appendage as a molecular reporter to investigate the differences between α3 and α5 chain variants. This study aims to elucidate the pathogenic effects of the Z-appendage in the α5 chain of collagen IV and deepen our understanding of its role in disease mechanisms.

## Results

### Generation of the *Col4a5*-Zurich variant knock-in mouse

To assess pathogenicity, we developed a mouse harboring the Zurich variant on the α5 chain of collagen IV using CRISPR/Cas9 editing technology. We used two guide RNAs and a 490-base single-stranded oligonucleotide for repair. For genotyping, we performed PCR amplification using COL7-COL8 primers and sequenced with COL7; all oligonucleotides are indicated in Figure 1A. As shown in Figure 1B, we introduced mutations substituting the last threonine with eight new amino acids: QQNCYFSS. We generated hemizygous males and homozygous females, then expanded and studied the line. Hemizygous male and homozygous female mice were viable, fertile, and born in the expected Mendelian ratios.

**Figure 1.**
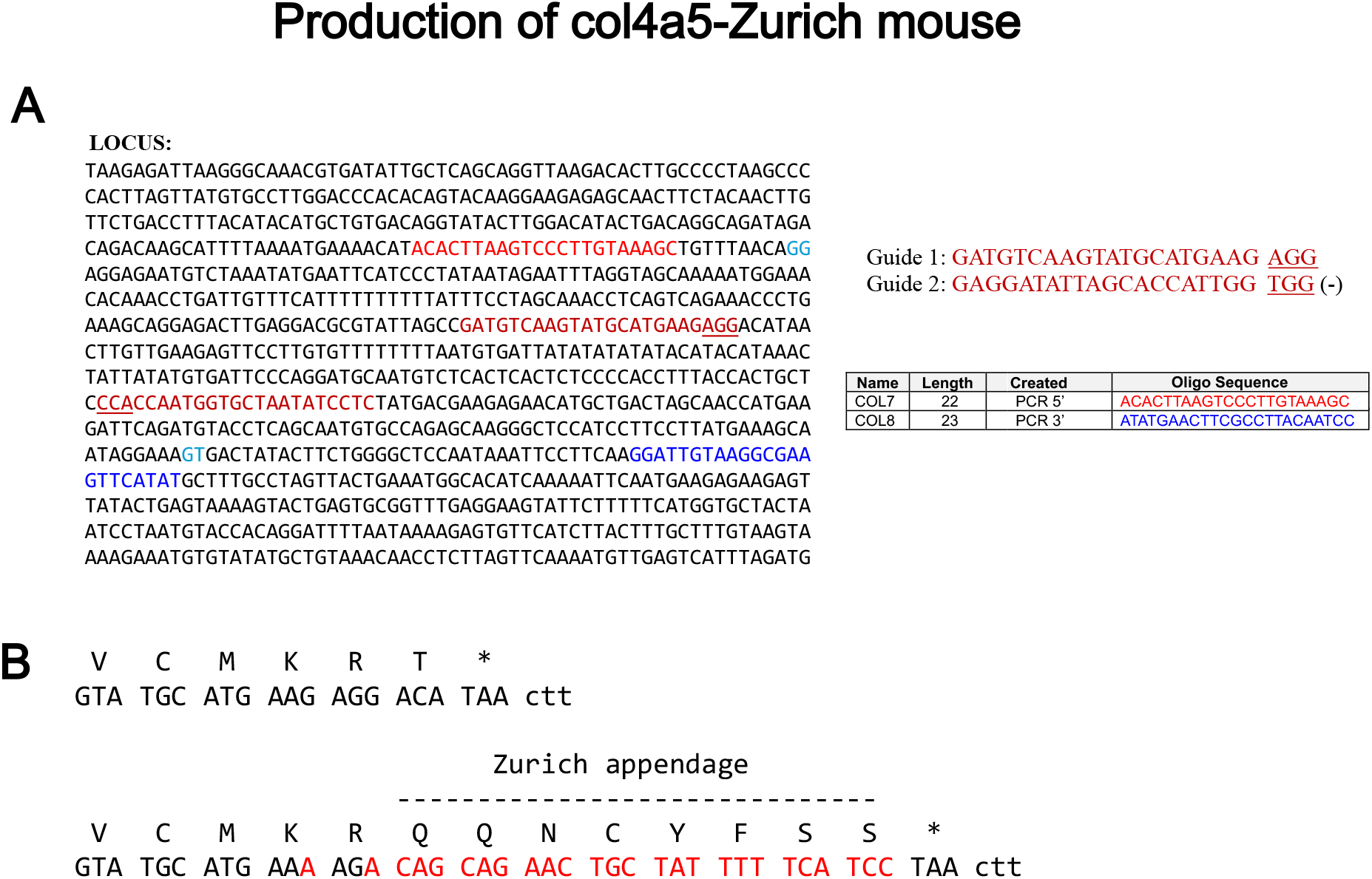
Production of *Col4a5*-Zurich mouse. The mice were created using CRIPR/Cas9 technology. (**A**)The sequence of the targeted locus is shown with the guide RNAs and genotyping primers highlighted with different colors. (**B**) shows the resulting amino acid addition at the end of the endogenous sequence of α5 chain of collagen IV.

### Collagen IV^α345^ scaffold is deposited in the kidneys of *Col4a5*-Zurich mice

We have analyzed the deposition of collagen IV ^**α345**^ scaffold to the GBM of *Col4a5*-Zurich mice. First, whole kidney homogenates from mutants and wild-type controls were probed for the presence of α3, α4, and α5 chains by western blot (Fig.2A). We found no difference between controls and *Col4a5-*Zurich homozygous female mouse samples. Hexamer formation and cross-linking were also unaffected.

**Figure 2.**
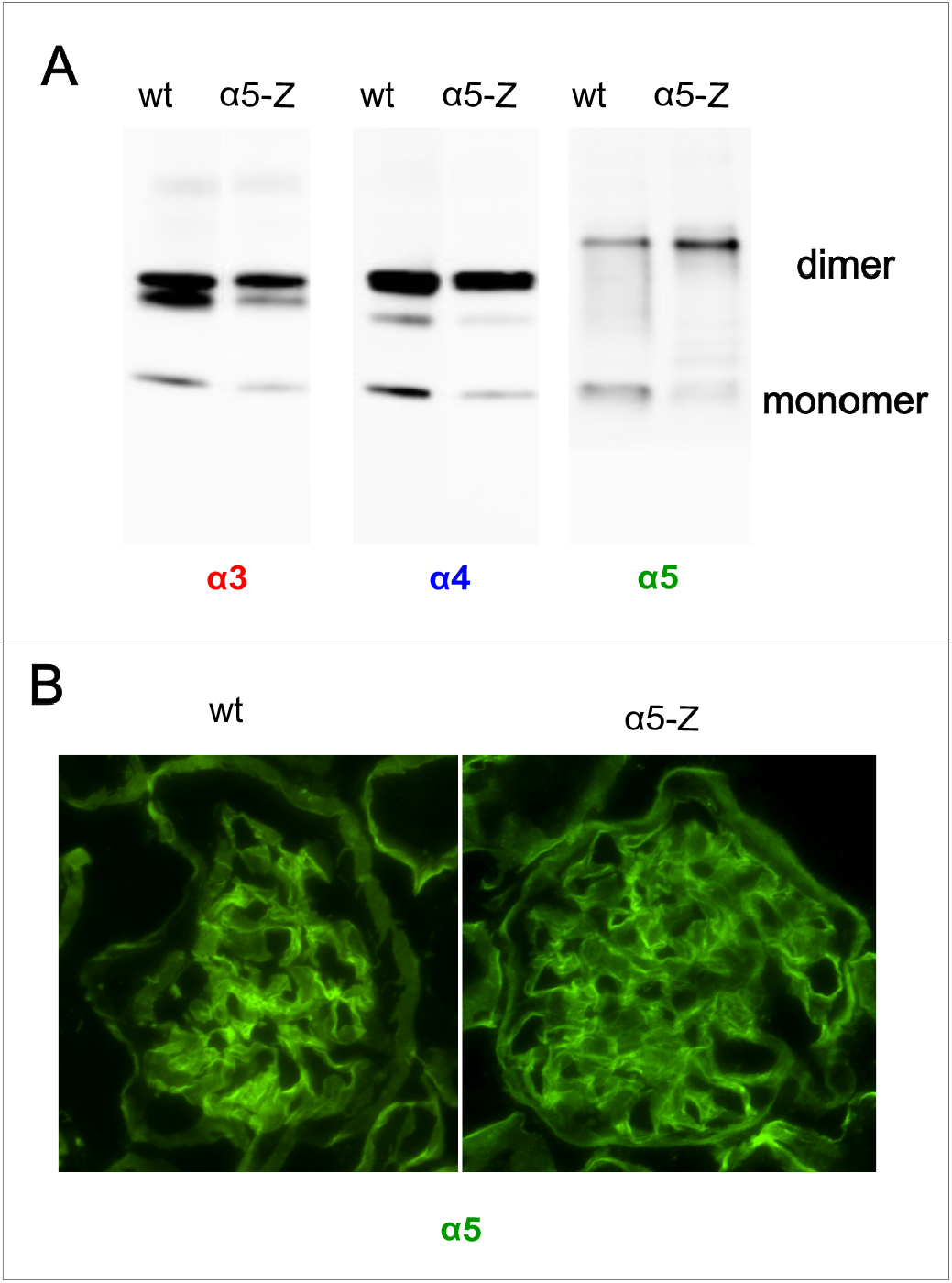
Collagen IV α5 harboring Z-appendage is incorporated into the GBM and Bowman’s capsule. (**A**) Western blot showing expression of α3, α4, and α5 chain of collagen IV in kidneys of homozygous female mutants and controls. The following chain specific antibodies were used: H31(α3), H43(α4) and Mab5(α5). The presence of Z-appendage did not alter expression of collagen chains and α345 hexamer formation in kidney. (**B**) Indirect immunofluorescence of the glomeruli of wt and mutant *Col4a5-Z* mice for α5 chain of collagen IV using chain specific antibody. In both wt control and mutant mice α5 antibody stained the GBM and Bowman’s capsule. This shows that mutant α5 chain carrying Z-appendage is well incorporated in the GBM.

Second, we performed immunofluorescent staining to localize the α5 chain of collagen IV in the kidney sections of mutants and wild types. The α5 chain was present in the GBM of the genetically modified animals at a level similar to that of the wild types (Fig.2B). Thus, Zurich appendage at the NC1 domain of collagen IV α5 chain does not cause a disruption of α345 scaffold assembly. This is in correlation with our findings for the Zurich appendage attached to the α3 chain of collagen IV [15].

### The kidney filtration function of *Col4a5*-Zurich mice is unaffected

We first qualitatively assessed proteinuria in female homozygous *Col4a5*-Zurich mice. As shown in Figure 3A, urine samples from 8-week-old and 28-week-old homozygous females were resolved on SDS-PAGE, along with samples from age-matched wild-type controls and reference mouse albumin. In the analyzed *Col4a5*-Zurich mutant females, no substantial albumin was detected in the urine (Fig. 3A). In contrast, moderate albuminuria was previously observed at comparable ages in *Col4a3*-Zurich mice [15]highlighting a difference between the two models.

**Figure 3.**
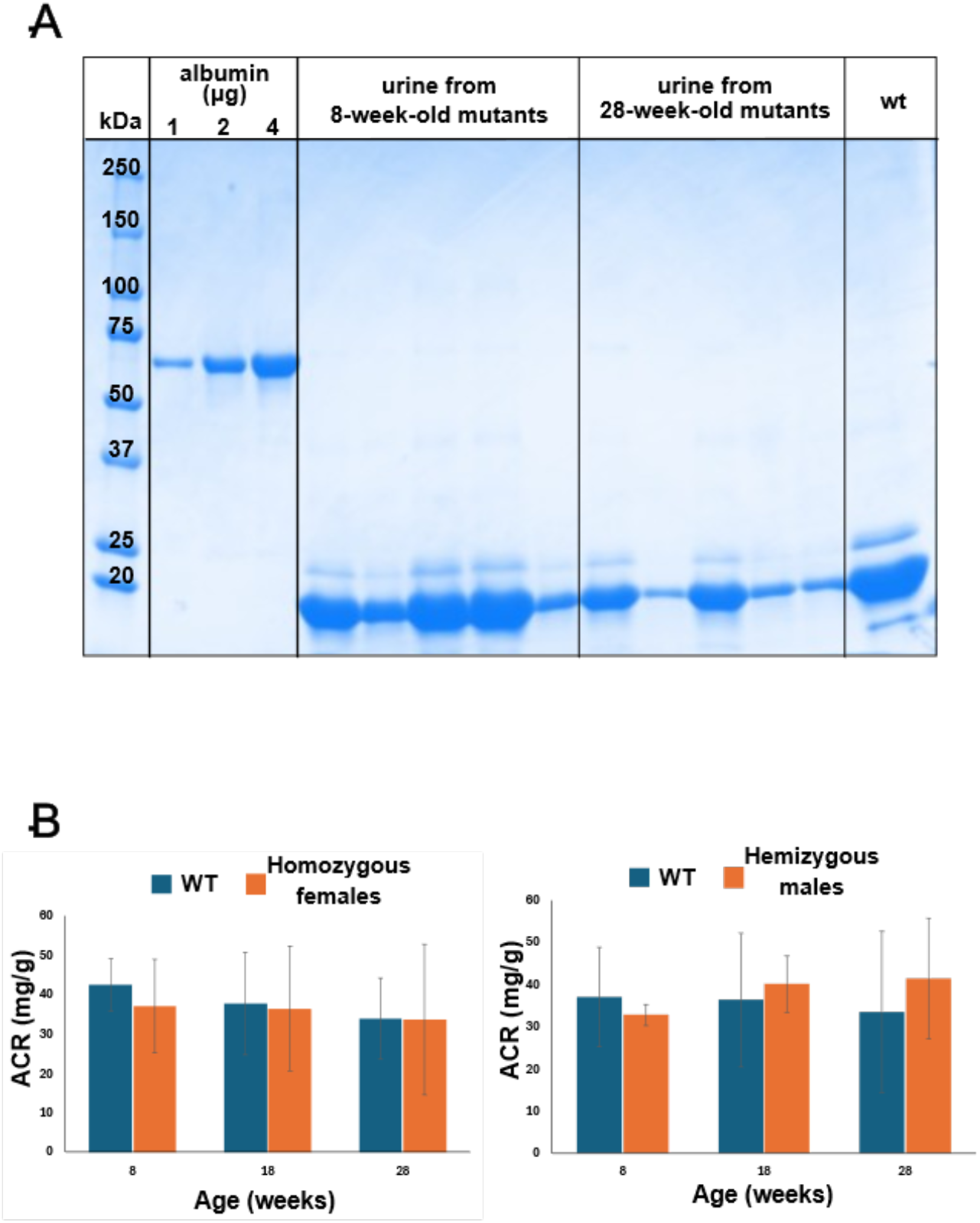
The *Col4a5-Z* mutants do not have albuminuria. **(A)** SDS PAGE of urine samples collected from homozygous *Col4a5*-Zurich mutants (n=5) at 8 weeks and 28 weeks. BSA used as standard. None of the mice at 28 weeks showed albuminuria.**(B)** Albumin to creatinine ratios (ACR) measured in urine of homozygous (n=5) and wild type (WT) control (n=2) mice at age 8, 18, and 28 weeks. No differences in the ACR values were observed at any age.

For the quantitative analysis, we measured albumin/creatinine ratios (ACRs) in the urine of controls, homozygous females, and hemizygous males at 8, 18, and 28 weeks of age (groups of 5 animals each). We found no differences in ACR values among these groups in both younger and older mice (Fig. 3B). Thus, while albumin crosses the glomerular filtration barrier in *Col4a3*-Zurich mice, it does not in *Col4a5*-Zurich mice.

### The Bowman’s capsule in *Col4a5*-Zurich mice is thickened with excessive deposition of collagen

Besides the absence of proteinuria in the *Col4a5*-Zurich mice, histological analysis of the 28 week-old hemizygous males and homozygous females showed occasional partial glomerular sclerosis (arrowhead in Fig. 4A) and, most notably, thickening of the Bowman’s capsule (arrows Fig. 4A). The PAS stains the glyco-residues; thus, the thickening of the Bowman’s capsule is likely due to the excessive extracellular matrix deposition. It is unclear; however, what molecules are deposited in the Bowman’s capsule of the mutants.

**Figure 4.**
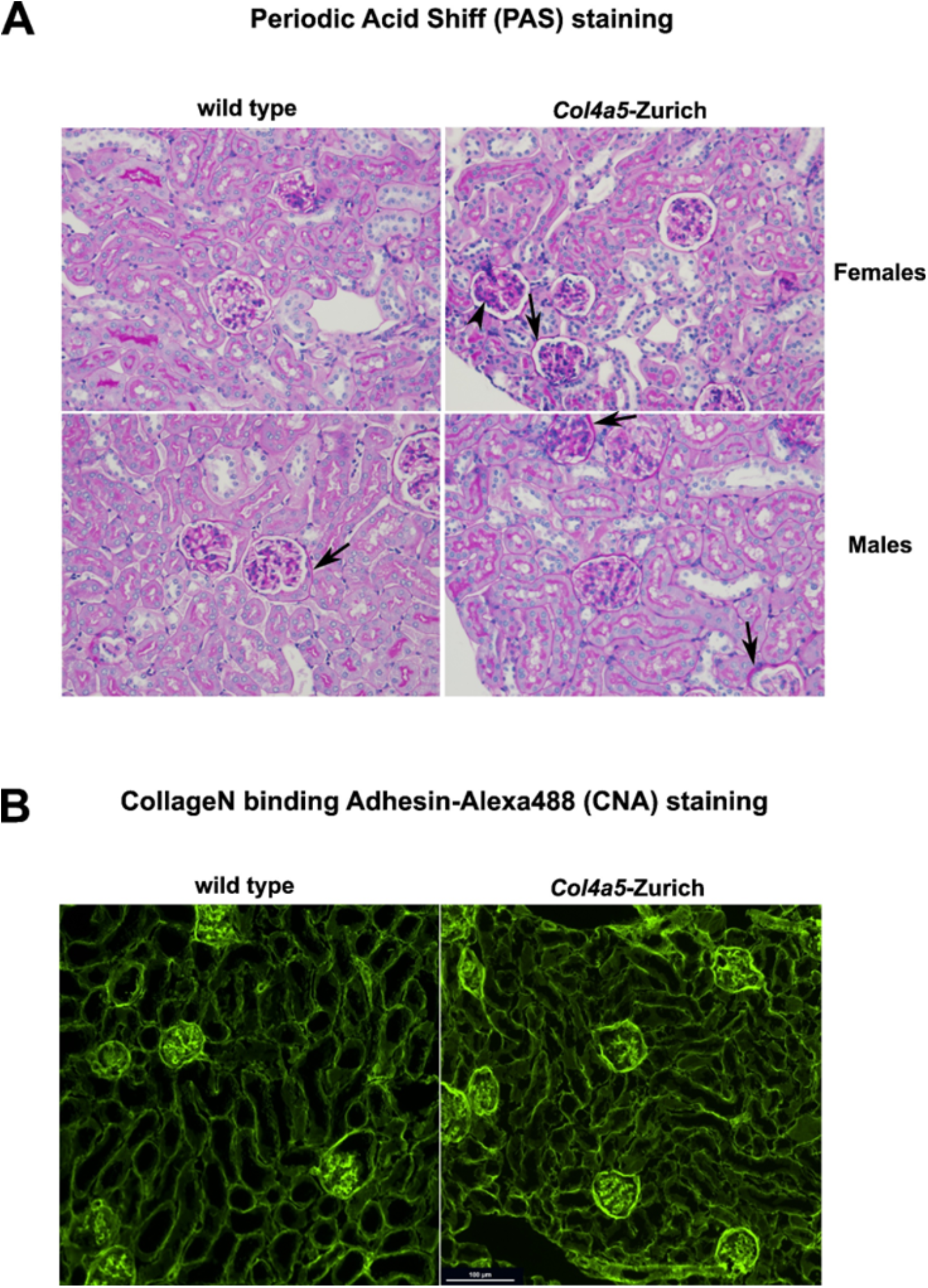
Histology and collagen staining of the *Col4a5-Z* kidney sections. **(A)** PAS histology staining of *Col4a5-Z* mutant and wild type control paraffin kidney sections. Arrows point to thickened Bowman’s capsule, arrowhead to a focal glomerular sclerosis. **(B)** Collagen is visualized in the frozen kidney sections with fluorescently labeled CNA35 trimer [18]Note increased staining of the Bowman’s capsule in mutants.

To identify contributing factors for the thickened Bowman’s capsule we first performed staining with the fluorescently labeled CNA35 trimer. The <u>C</u>ollage<u>N</u> binding <u>A</u>dhesin (CNA) protein from *Staphylococcus aureus* has been used to bind collagen’s triple helices in tissue sections and blots. We also recently reported an improved collagen affinity for the CNA35 trimer [18]. When stained with the CNA35 trimer, *Col4a5*-Zurich mutant mice show substantial thickening of the Bowman’s capsule compared to the wild type controls (Fig. 4B). Therefore, there is excessive deposition of collagen in the Bowman’s capsule of *Col4a5*-Zurich mutants.

### Composition of basement membrane in the Bowman’s capsule of *Col4a5*-Zurich mice is shifted towards higher content of collagen IV^α121^

To further analyze Bowman’s capsule composition in the mutants, we compared immunofluorescent staining for α1 and α5 collagen IV chains. As seen in Figure 5, the amount of α1 chain in the Bowman’s capsule of *Col4a5*-Zurich mice is increased compared to the wild types. At the same time, there is similar intensity of α5-specific staining in the Bowman’s capsule of the mutants in comparison to the controls (Fig. 2B). Therefore, the composition of the basement membrane in the Bowman’s capsule of the mutants is shifted towards increased collagen IV ^**α121**^ scaffold.

**Figure 5.**
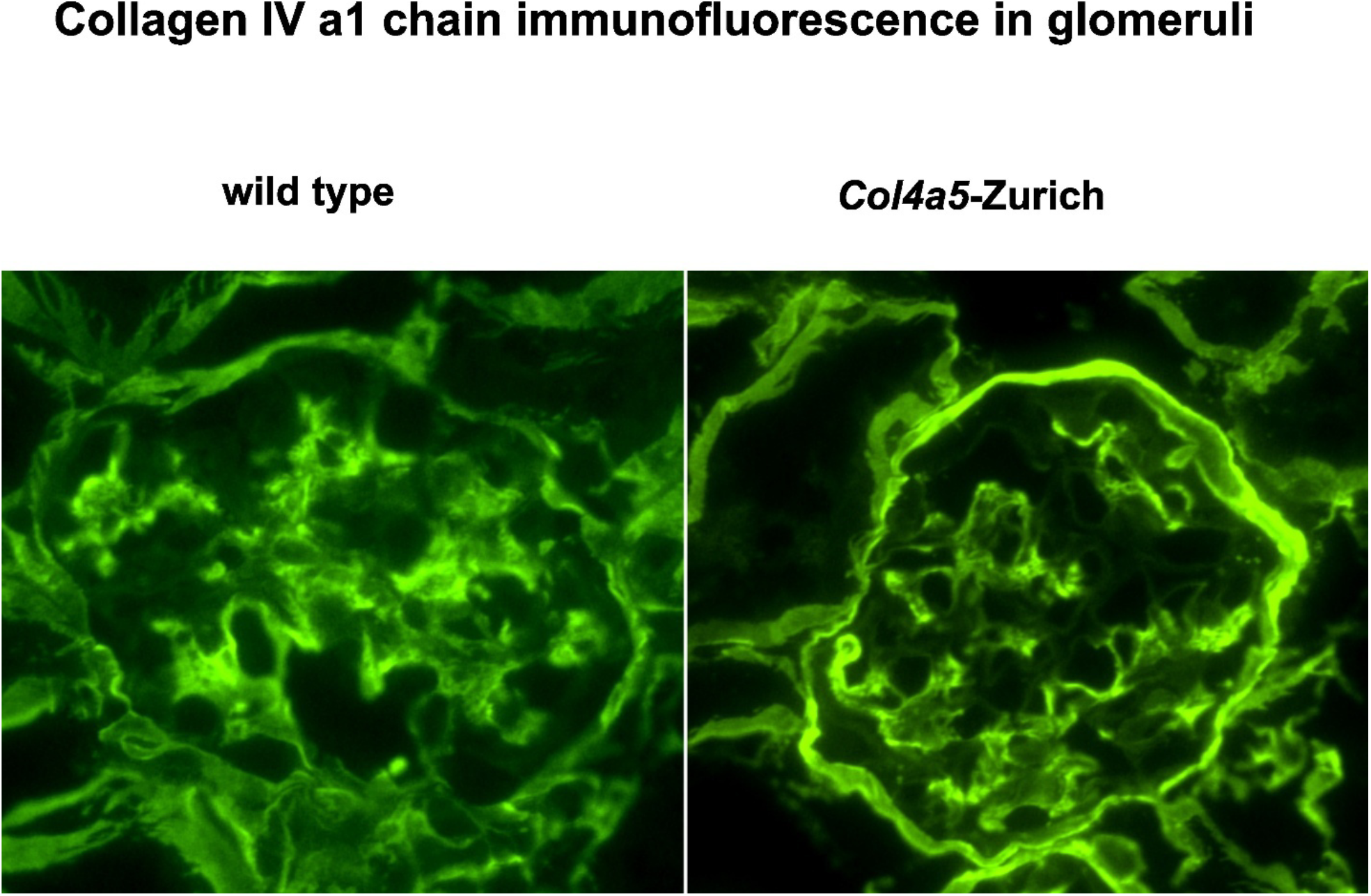
Immunofluorescent staining of the α1 chain of collagen IV in *Col4a5-Z* and wild type kidney sections. Immunofluorescence shows increased content of the α1 chain of collagen IV in the mutant Bowman’s capsule.

### Electron microscopy shows minor alterations in the GBM and morphological defects in the Bowman’s capsule of the *Col4a5*-Zurich mutants

To explore the kidney ultrastructure of the *Col4a5*-Zurich mice we employed transmission electron microscopy (TEM). The ultrastructural morphology of the glomerular filtration barrier in the 28 weeks old homozygous female was mostly normal. However, the glomerular basement membrane at some places was irregular in thickness with slight occasional splitting (Fig. 6A, arrowhead) present. Most notably, the basement membrane of the Bowman’s capsule was significantly thickened and deposition of the fibrillar collagens surrounding basement membrane was observed (Fig. 6B). In addition, the Bowman’s capsule is thickened with the cellular component (cell is indicated in Fig. 6B).

**Figure 6.**
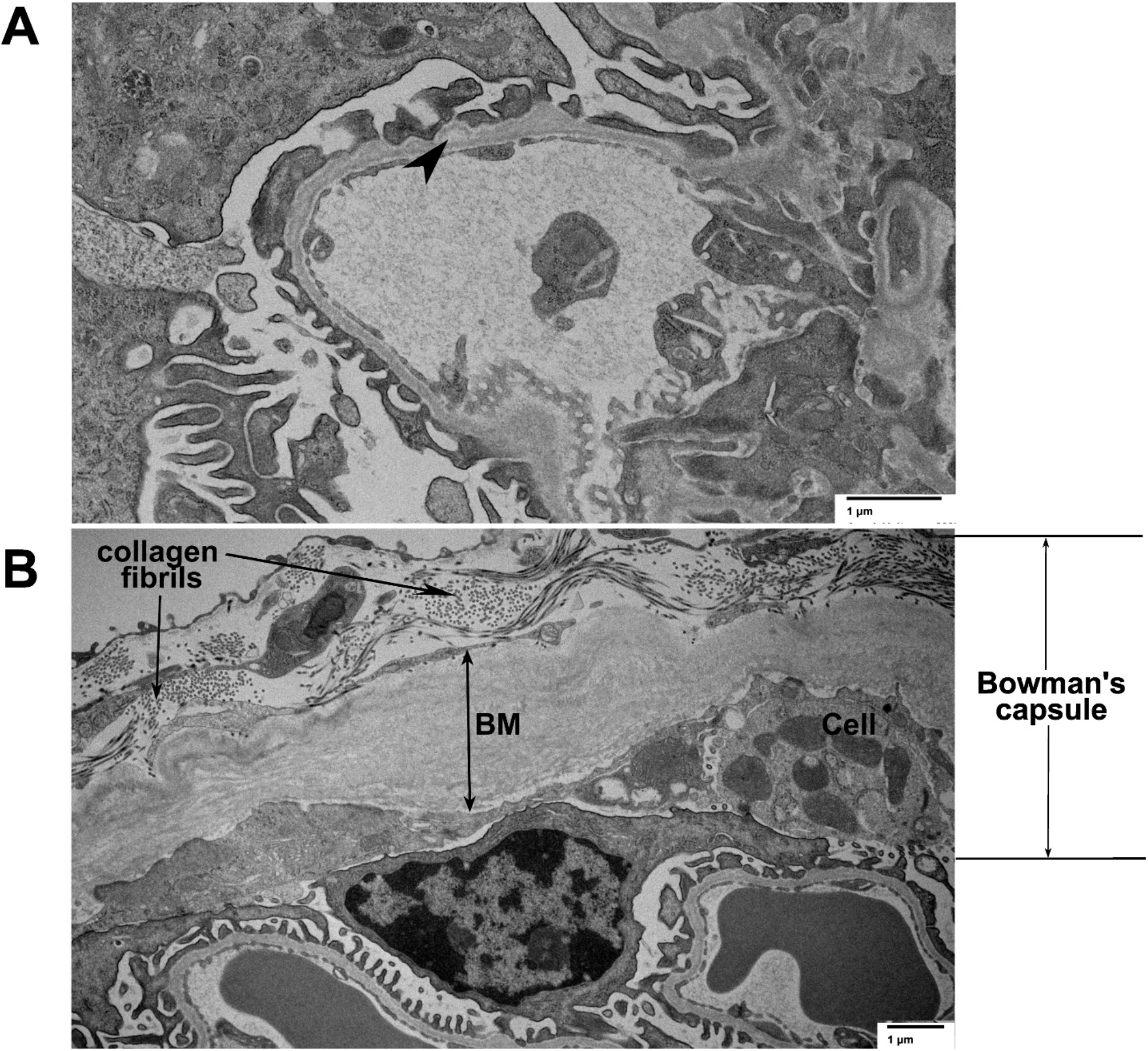
Transmission electron microscopy of the *Col4a5-Z* mutant kidney. **(A)** Capillary loop in the glomerulus of the 28-week-old *Col4a5-Z* mutant. The filtration barrier, including podocytes and glomerular basement membrane, are generally well preserved with minor occasional splitting of the GBM (arrowhead). **(B)** Bowman’s capsule is shown. The basement membrane (BM) is thickened, deposits of collagen fibrils are observed as well as cellular deposits (Cell).

## Discussion

Alport syndrome (AS) is caused by genetic variants that lead to defects in the collagen IV scaffold [4]. Collagen IV is a trimeric molecule essential for the structure of basement membranes in all tissues. Its composition is organ-specific, with the α345 isoform predominating in the glomerular basement membrane (GBM) and the α565 isoform in Bowman’s capsule [1].

In this study, we developed and analyzed a *Col4a5-Zurich* mouse model in which an eight-amino-acid appendage is attached to the C-terminal non-collagenous domain (NC1) of the collagen IV α5 chain. Previously, the same appendage on the α3 chain of collagen IV was found to cause AS [15]. Our goal was to better understand genotype-phenotype correlations in AS, particularly X-linked AS, by investigating the *Col4a5-Zurich* mice. Unlike the *Col4a3-Zurich* mice, the new model exhibits no albuminuria. Similar to the α3-Z mutants, the α5-Z mice assemble and deposit the collagen IV^**α345**^ scaffold into the GBM. While minor morphological alterations of the GBM are observed in older α5-Z mutants, these changes are insufficient to compromise the filtration barrier. Interestingly, we found the most pronounced defects in Bowman’s capsule of the *Col4a5-Zurich* mice.

The predominant isoform of the collagen IV scaffold in the GBM is α345, while the α565 isoform dominates in the basement membrane of Bowman’s capsule. The structure of the NC1 hexamer involving α565 in mutant mice may be more vulnerable due to the presence of two α5 chains carrying the Z-appendage. Immunofluorescence data revealed that the mutant α5 chain is incorporated into Bowman’s capsule (Fig. 2B), but there is increased deposition of the α1 chain compared to controls (Fig. 5B). This suggests that defects in the collagen IV ^**α565**^ protomer may be partially compensated by increased α121 deposition.

A recent study of a mouse model of X-linked AS, harboring a human nonsense mutation in *Col4a5*, demonstrated that crescent formation is the primary driver of disease progression. These crescents arise from damage to both the filtration barrier (GBM) and the capsular barrier (Bowman’s capsule) [13]. In addition, two recent case studies of X-linked AS with variants in the α5 chain were reported that have attracted our attention with regard to our mouse model. Both suggested that defects in collagen IV scaffold assembly begin in Bowman’s capsule before affecting the GBM [19], [20]. Interestingly, in both cases the variants were Gly to Ser substitutions within the triple helical region close to the NC1 domain.

To investigate the structural impact of the Z-appendage on collagen IV NC1 hexamers, we performed structural predictions in different contexts. In the α345 hexamer, the Z-appendage on the α5 subunit caused minimal impact on the predicted structure (Fig. 7A). However, an unexpected combination of secondary structural features arose in the presence of the Z-appendage on the collagen IV ^**565-121**^ hexamer (Fig. 7B). The presence of two Z-appendages, positioned in close proximity, resulted in the formation of a novel structure at the junction where the triple helix connects to the NC1 domain. This structure includes a three-stranded antiparallel beta sheet: the first strand (“1” in Fig. 7B) is formed by the “TSSV” sequence in the triple helix of one α5 subunit, while the other two strands (“2” and “3” in Fig. 7B) are derived from the Z-appendages. This structure may be further stabilized by a disulfide bond, as the cysteine residues of the Z-appendages are predicted to be positioned close enough to form such a bond. The increased rigidity induced by the described secondary structure may negatively impact the packaging, transport of α565 promoters, and the overall integrity of the assembled scaffold. Additionally, alterations near the junction of the NC1 domain and triple helix may potentially hinder the interactions with other basement membrane proteins or cell surface receptors.

**Figure 7.**
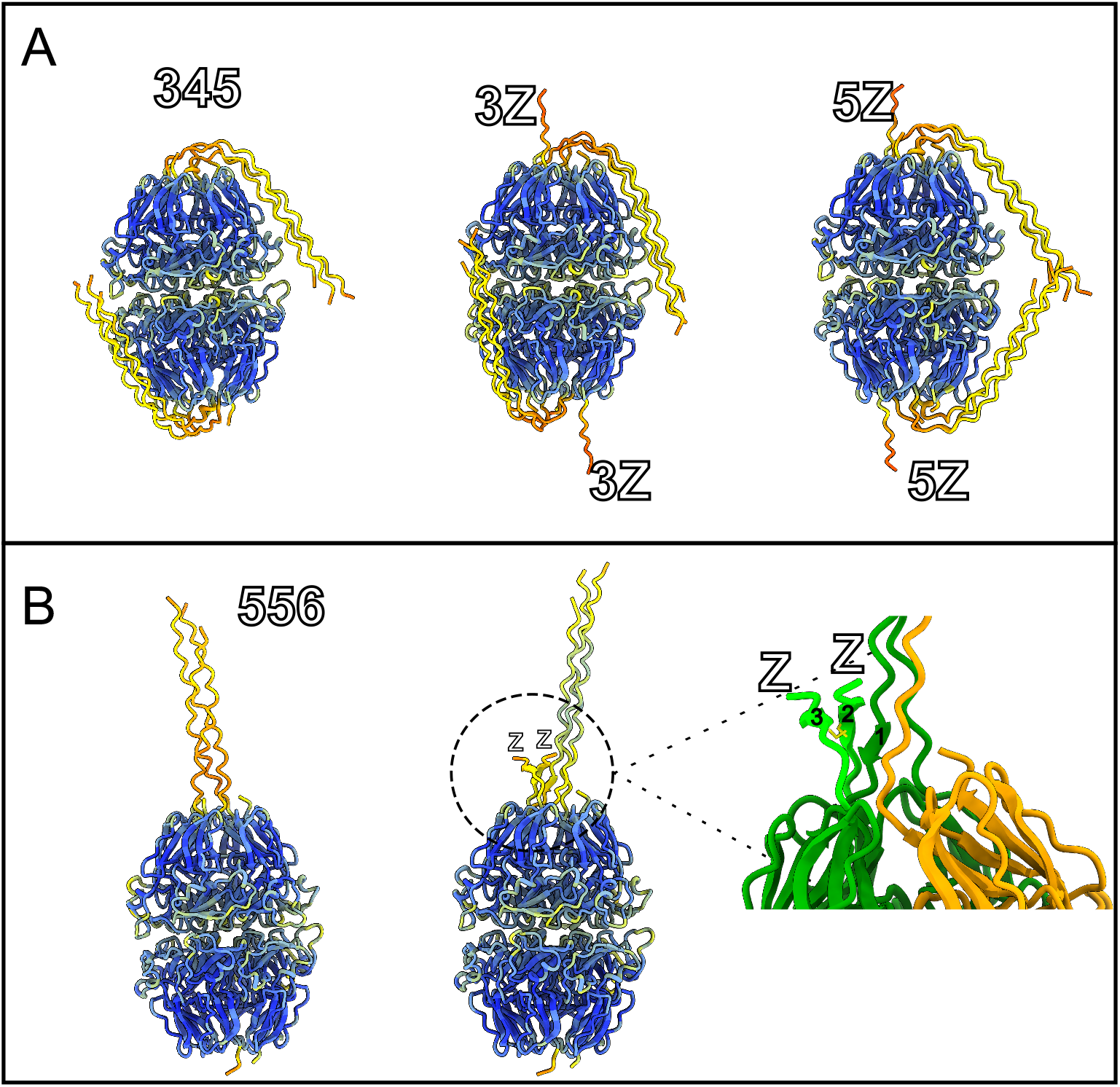
3D modeling of the collagen IV hexamers of different compositions. **(A**) AlphaFold3 prediction of collagen IV ^**α345**^ hexamer, and α345 hexamer with Z-appendage on the α3 and α5 subunits colored by pLDDT. (**B)** AlphaFold3 prediction of collagen IV ^**α565- α121**^ hexamer, and α565-α121 hexamer with Z-appendages on the α5 subunits colored by pLDDT. On the right is magnified region of AlphaFold 3 prediction collagen IV ^**α565- α121**^ hexamer with Z-appendages on the α5 subunits with α5 colored in green, Z-appendage in light green, α6 in yellow, and cystine residue sidechains in the z appendage shown in yellow. “1”, “2”, and “3” refers to the β-sheet structure formed by triple helix and Z-appendages.

This study provides a direct comparison between two mouse models with identical variations in the NC1 domain of the α3 and α5 chains of collagen IV. Surprisingly, the phenotypes differ significantly: while *Col4a3-Zurich* mice develop slow-progressing AS, *Col4a5-Zurich* mice show no functional abnormalities in the filtration barrier. However, defects in Bowman’s capsule in the *Col4a5-Zurich* mice highlight the functional importance of the collagen IV ^**α565- α121**^ scaffold. Furthermore, our structural modeling points toward specific bioactivity of the triple helical/ NC1 domain junction. To our knowledge, this is the first study demonstrating that the α565 scaffold has a unique function.

## Experimental Procedures

### Isolation of NC1 domain from mouse kidney for Western Blotting

The wild-type control and homozygous mouse kidney were finely chopped over ice and resuspended in ice cold TBS buffer containing 0.1% Tween 20 (TBST) supplemented with protease inhibitors. Mixture was centrifuged at 4000 xg for 5 mins at 4 C and pellet was collected. Pellet was washed 4 times with TBST. Final collected pellet was resuspended in 250 ul of collagenase digestion buffer[21] containing 100 ug/ml final concentration of collagenase followed by shaking and overnight incubation at 37 C. Supernatant was collected by centrifugation at 6000 xg and separated in 4-20% gradient gel using SDS-PAGE and transferred to nitrocellulose membrane using Trans-Blot semidry transfer cell (BioRad). Transferred membranes were probed with H31, H43 and Mab 5 antibodies for the detection of alpha 3/4/5 chains of collagen IV respectively.

### Mouse Urine Collection for ACR Measurements and SDS gel analysis

Urine at different age of experimental mice were collected following the animal protocol#M1900063-01 approved by the MC IACUC. Urine albumin concentration were quantified using the Albuwell M kit (Exocell, Inc) kit and analyzed using online ELISA calculator from Arigo Biolaboratories (https://www.arigobio.com/elisa-calculator). Urine creatinine concentration were assessed using creatinine (enzymatic) reagent set (Cat#10022-480) from Pointe Scientific, Inc. in accordance with the manufacturer’s protocol. Additionally, SDS-PAGE electrophoresis was conducted on 2 μl of urine for majority of samples using Biorad 4-20% gel.

### Transmission electron microscopy of mouse kidney

Freshly isolated kidneys were sliced into small pieces (2 × 2 mm) and fixed overnight in 2.5% glutaraldehyde buffered with 0.1 M sodium cacodylate buffer, pH 7.5. The samples were post-fixed in 1% osmium tetroxide, followed by dehydration through a graded ethanol series to 100%. After dehydration in propylene oxide, the samples were infiltrated and embedded in Spurr’s epoxy resin. Ultrathin sections (70 nm) were placed on 300 mesh copper grids and stained with 2% uranyl acetate and Reynold’s lead citrate. The stained sections were observed using a T-12 electron microscope (Philips/FEI) at 100 kV, and images were captured using a 2K camera (AMT).

### Immunofluorescence of Mouse kidney sections

The kidney sections were snap frozen in dry ice-ethanol slurry using OCT compound (Tissue-Tek). Slides of 5 μm kidney sections were prepared after cryosectioning. Slides were dried at room temperature for 10 mins, washed with PBS and fixed in ice cold acetone for 10 mins and washed with PBS. For CNA staining, slides were blocked with 10% goat serum for 10 mins and 10 mins in 1:1000 dilution of CNA in 1% goat serum. Image was taken after washing slides with PBS 3 times. For chain specific collagen IV antibodies incubation, slides were pretreated with 6M urea in 0.1M glycine pH 3.5 for 10 mins to uncover the antigen, washed with PBS and PBS with 0.2% Tween. Slides were then pre-incubated with 10% normal goat serum (Invitrogen) for 1 hour at room temperature (RT) to block non-specific binding of antibody and incubated overnight in humidified chamber at 4 °C with rat anti-collagen IV α1 NC1 (1:250 dilution, H11), rat anti-collagen IV α3 NC1 (1:250 dilution, H31), and rabbit anti-collagen IV α5 NC1 (1:250 dilution, Col4a5 Ab). The H11 and H31 antibodies were kindly provided by Y. Sado (Shigei Medical Research Institute) while ColIVa5 Ab is from Bicell scientific (cat# 05005). Secondary antibodies used for immunofluorescence were Alexa488 goat anti-rat and goat anti-rabbit (1:1000 dilution; Abcam) and incubated for 1 hr at room temperature. Antibodies were diluted in PBS/0.1% Tween and 5% normal goat serum and slides were washed with PBS with 0.2% Tween. A negative control slide eliminating primary antibody incubation step was also prepared in a similar way.

## Notes

### Competing Interest Statement

The authors have declared no competing interest.

### Funding

- National Institutes of Health: R01DK131101, R01DK018381

